# Saracatinib and Dasatinib Fail To Prevent Heritable Pulmonary Arterial Hypertension

**DOI:** 10.1101/345447

**Authors:** Reid W. D’Amico, Santhi Gladson, Sheila Shay, Courtney Copeland, James D. West

**Affiliations:** Department of Biomedical Engineering, Vanderbilt University, Nashville, Tennessee, 37232, USA; Division of Allergy, Pulmonary, and Critical Care Medicine, Vanderbilt University Medical Center, Nashville, Tennessee, 37232, USA.

**Keywords:** Pulmonary Hypertension, SRC

## Abstract

Evidence suggests that the deregulation of SRC Family Kinases may play a role in the development of heritable pulmonary arterial hypertension, associated with BMPR2 mutations. The truncated c-terminus of the BMPR2 protein is known to increase the phosphorylation and downstream activity of SRC tyrosine kinases. To test the hypothesis that the inhibition of SRC can prevent heritable PAH due to a BMPR2 mutation, we exposed BMPR2 mutant mice to SRC inhibitors, saracatinib and dasatinib, to block the SRC activation caused by the BMPR2 mutation. Saracatinib and dasatinib failed to prevent the development of PAH in BMPR2 mutant mice. Increased pressure in the right ventricle was not normalized and muscularization of large blood vessels was not reduced when compared to wild type mice. Inhibiting SRC’s phosphorylation does not prevent heritable PAH, and thus supports evidence that SRC’s aberrant localization and trafficking in PAH plays a more critical role in disease development.

## INTRODUCTION

Pulmonary arterial hypertension (PAH) is a progressive and fatal illness of the pulmonary microvasculature. The remodeling of the small resistance arteries in the lung leads to increased pulmonary vascular resistance, increasing the workload on the heart’s right ventricle (RV) until it eventually fails (1, 2).

The most studied heritable risk factor for the development of PAH is a mutation in the type 2 receptor in the BMP pathway (BMPR2). However, the mechanism underlying BMPR2 and the development of PAH is not well understood (3-6). The structural deformity seen in the mutated BMPR2 protein is known to play a role in a host of deregulated protein pathways in PAH. In heritable BMPR2 mutations, the c-terminus of the protein is truncated. This results in the increased phosphorylation of proteins responsible for maintenance of cellular mechanisms like growth, ECM maintenance, and cell division (3,7). Abnormalities in cell-cell and cell-ECM force transduction contribute to the mechanical pathology seen in the resistance vessels (8). When considering the abnormal mechanical properties of the vessels in the lung, it is important to probe the molecular mediators of ECM regulation, mechanotransduction, and intercellular force transduction, as many of these mediators represent potential therapeutic targets (9).

Over recent years, considerable evidence has been accumulated suggesting the involvement of receptor tyrosine kinases (RTK) in the pathogenesis of pulmonary vascular remodeling seen in PAH. SRC Family Kinases (SFKs) are the largest subfamily of non-RTKs consisting of 9 kinases, known as SRC tyrosine kinase or SRC, which share similar structures and function (10). SFKs play an important role in regulating signals from many RTKs and have evolved many complex regulatory strategies that couple with the cytoplasmic domains of many proteins. This complex regulation by SFKs is what controls many signaling pathways required for DNA synthesis, control of receptor turnover, differentiation, actin cytoskeleton rearrangements, motility, and survival (11,12). The relationship among SFKs, vascular remodeling, and the pathogenesis of PAH are not well-explored.

In heritable PAH due to BMPR2 mutations, SRC’s phosphorylation and downstream activity is increased (3,7). SRC binds to the cytoplasmic tail of the mutated BMPR2 protein (13), which is likely a key component in the development of PAH (14). We hypothesize that inhibiting SRC’s phosphorylation will prevent its downstream activities and may prevent the development of heritable PAH. To test this hypothesis, we studied the ability of two known molecules that prevent SRC’s phosphorylation, saracatinib and dasatinib (15, 16).

## MATERIALS and METHODS

### Heritable PAH: BMPR2 Mutant Mice

When exposed to doxycycline, Rosa26-Bmpr2^R899X^ mice will express the patient-derived R899X mutation in all tissues. After transgene activation, about 50% of Rosa26-Bmpr2^R899X^ adult mice (10-14 weeks of age) will develop PAH as characterized by right ventricular systolic pressures (RVSP) above normal range (3). BMPR2 mutant mice were fed a western diet consisting of doxycycline (Bioserv) at 0.2g/kg for 6 weeks. Two weeks after the start of the diet, osmotic pumps (Alzet 1004) containing either dasatinib or saracatinib in 50% DMSO/50% 16α-hydroxyestrone (16-OHE) solution or vehicle with the same DMSO/16OHE formulation were implanted. 16-OHE was used to further drive the disease progression (18). Dasatinib, saracatinib, or vehicle were delivered at 1 mg/kg/day for the remaining four weeks. After completion, the mice were placed under surgical anesthesia (Avertin) and the RVSP was measured by inserting a catheter into the right heart through the right jugular vein in a closed-chested procedure, as previously described (17). Ten-second segments of RVSP measurement were extracted from the RVSP waveform, and the maximum RVSP was calculated by measuring the difference between the peak and trough of the wave.

After sacrifice, tissues were collected for further analysis. All surgical procedures were approved by the Vanderbilt institutional animal care and use committee (IACUC).

### Histology & Western Blots

After RVSP measurement, the lungs were flushed with 5 mL phosphate buffer solution (PBS) via perfusion through the right ventricle. To remove the blood, a small cut was placed in the left atrium. The lungs were then inflated with 0.8% low melt agarose.

The lungs from mice with or without an activated R899X mutation in the BMPR2 receptor were isolated, embedded with Optimal Cutting Temperature compound, and sectioned after the mice were treated for four weeks with saracatinib, dasatinib, or vehicle.

Lung sections were stained with α smooth muscle actin (αSMA, Sigma) and DAPI. To quantify αSMA positive vessels, an observer blinded to treatment group counted the numbers of fully muscularized vessels per field in 10 random 10x fields in three mice per group. αSMA positive vessels were stratified by diameter (<25um or 50-100um). Vessels were identified with a Nikon Eclipse Ti microscope and were measured along the longest axis. The apexes and edges of the lungs were avoided in histological quantification and observation.

SRC activity was quantified by measuring a known downstream protein activated by SRC, p-CAS and CAS. Antibodies used for Western blots were: p-CAS (Cell Signaling #4015, 1:1000). All phosphorylation proteins were normalized to their respective total protein (i.e. pCAS/CAS).

### Statistical methods

Statistics were performed using two factor ANOVA (+/- BMPR2 mutation, +/- SRC Inhibitor), with Fisher’s exact test t for comparisons between experimental groups. Statistics were performed within R.

## Results

### SRC Phosphorylation Inhibition Fails to Prevent PAH in BMPR2 Mutant Mice

ROSA26-rtTA x TetO7-BMPR2^R899X^ mice express a dominant negative form of BMPR2, the patient-derived R899X mutation when induced by doxycycline. This allows strong suppression of BMPR2 in adult mice while avoiding developmental defects associated with its suppression in development. These ROSA26-BMPR2^R899X^ mice spontaneously develop pulmonary hypertension with reduced penetrance when their transgene is activated for six weeks, associated with multiple molecular abnormalities including increased SRC phosphorylation.

To determine whether inhibition of this SRC activity would prevent pulmonary hypertension, age and sex matched wild type and ROSA26-BMPR2^R899X^ mutant mice were treated with saracatinib and dasatinib for the last four weeks during six weeks of transgene activation. Saracatinib and dasatinib were shown to inhibit the action of downstream targets in the SRC pathways (**Figure 1**), indicating proper osmotic delivery through the subcutaneously placed pumps to achieve direct SRC inhibition.

**Figure 1:**
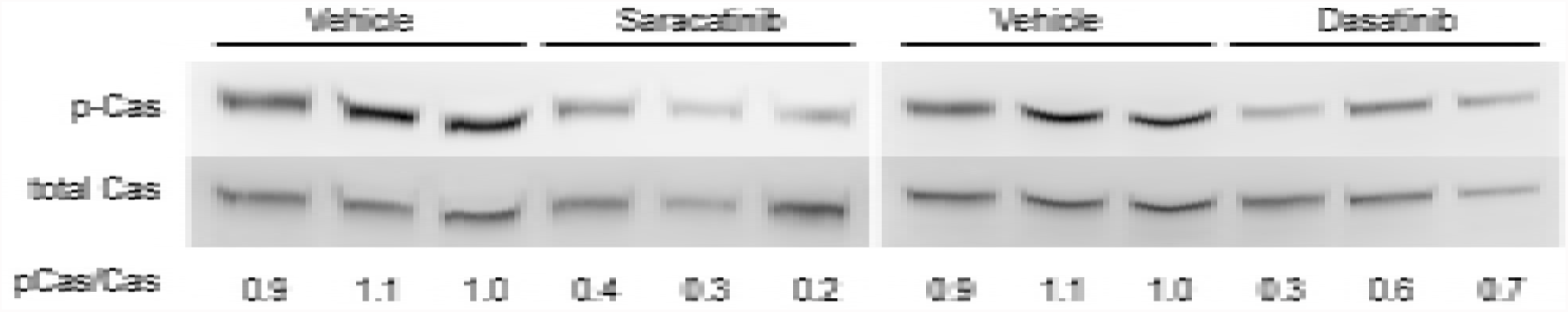
Saracatinib and Dasatinib reduce SRC phosphorylation and downstream activity. Western blots from ROSA26-BMPR2^R899X^mutant lung treated with saracatinib, dasatinib, or vehicle. Mutants have increased phosphorylation of SRC target, CAS. CAS phosphorylation and activity is reduced with saracatinib and dasatinib treatment.

While vehicle-treated mice developed elevated RVSP at about 35% penetrance, mice treated with SRC inhibitors have pressures indistinguishable from the vehicle-treated BMPR2 mutants (**Figure 2**). BMPR2 mutant mice have greater RVSPs than WT mice (p<0.05) and delivery of saracatinib and dasatinib to mutant mice does not reduce RVSP when compared vehicle treated mutants (p>0.05).

**Figure 2:**
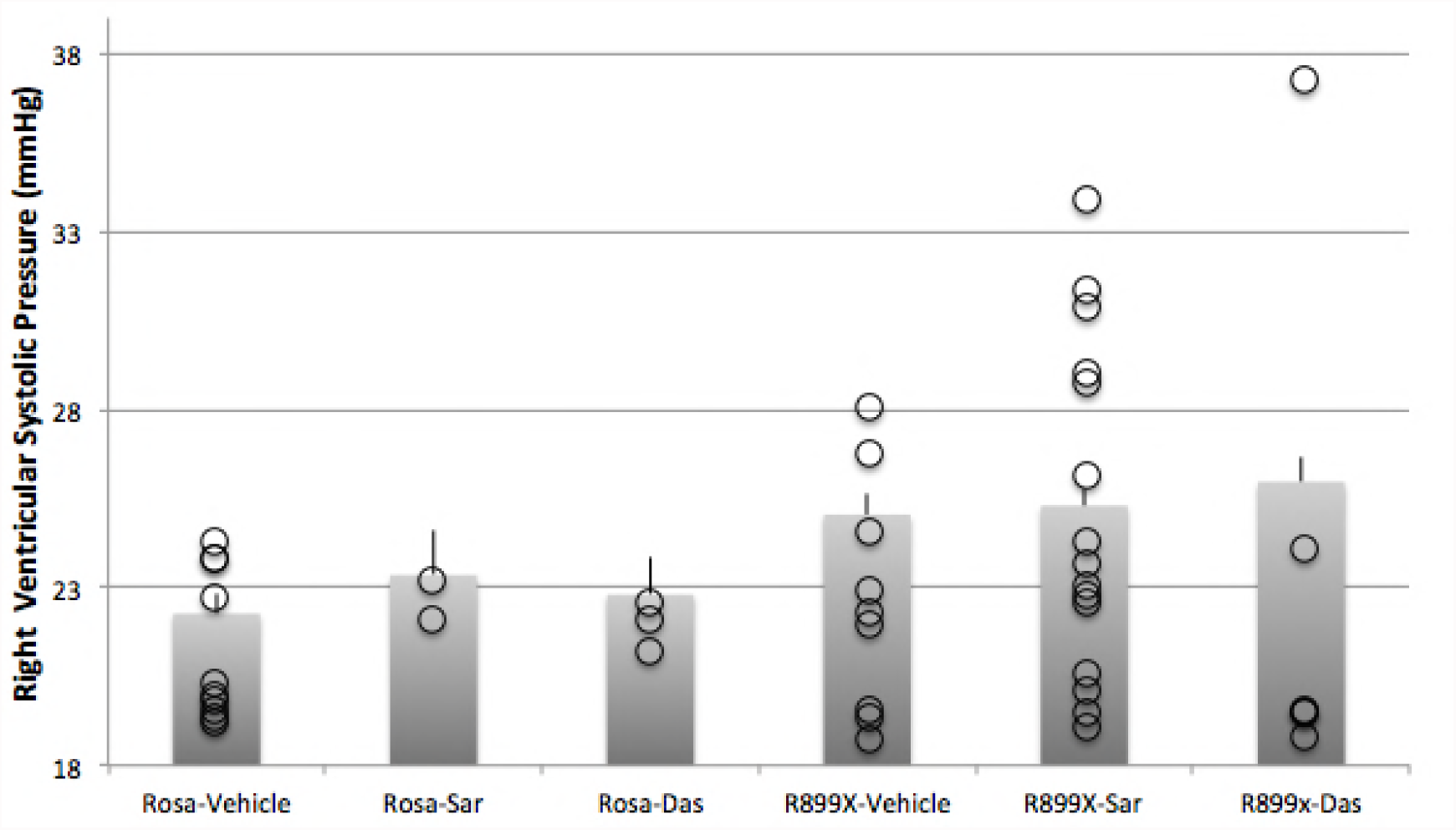
Saracatinib and Dasatinib do not lower RVSP in mutant mice. Right ventricular systolic pressures were significantly elevated (p<.001) in ROSA26-BMPR2^R899X^ mutants after six weeks of transgene activation through a western diet consisting of 1g/kg doxycycline. This elevation was not ameliorated through the administration of saracatinib and dasatinib pumps for the final four weeks. The circles represent the individual pressures of the mice and the bars represent the averages of all mice within the group. The error bars are reported as SEM.

Lung sections from ROSA26-BMPR2^R899X^ mutant mice had an increased number of large muscularized vessels when compared to the WT mice (p<0.05) (**Figure 3)**. The number of large muscularized vessels in the mutant mice was not reduced to the wild type mice by saracatinib and dasatinib treatment (p>0.05). The number of fully muscularized vessels is about doubled in all BMPR2 mutant mice, independent of treatment. All vessels were consistently measured along the longest axis and included the border of the masculinized vessels. Fully muscularized vessels were measured and vessels that did not have at least 75% vessel perimeter were excluded from the analysis. The muscles were stratified within Nikon and analyzed within R. All imaging channels were taken manually and overlaid in ImageJ.

**Figure 3:**
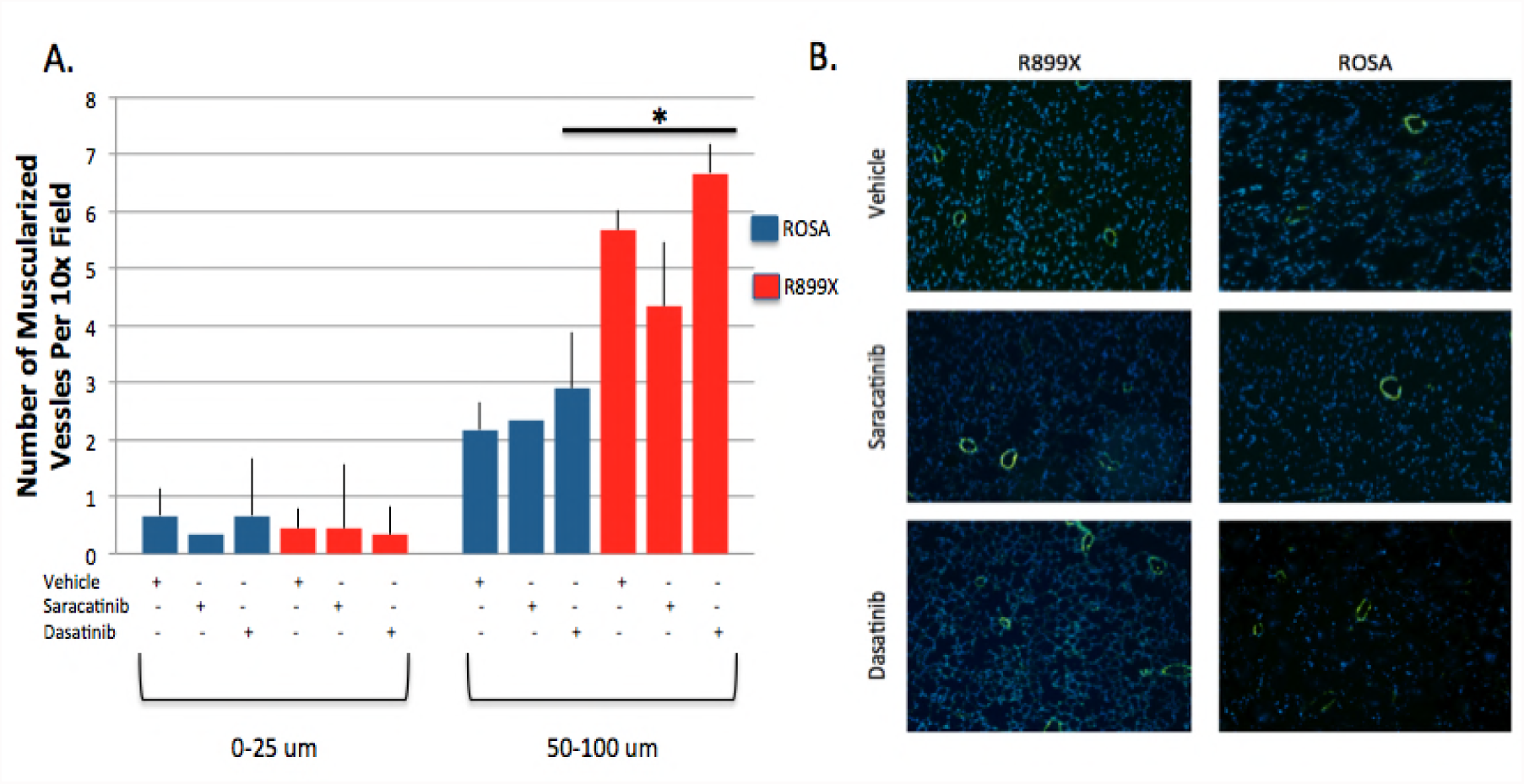
Saracatinib and Dasatinib do not normalize large muscularized vessels in mutant mice. **A)** ROSA26-BMPR2^R899X^mutants and wilt type mice have about the same number of small muscularized vessels per field. However, ROSA26-BMPR2^R899X^ mutants have about twice as many fully muscularized large sized vessels (50-100um). Saracatinib and dasatinib administration did not normalize the number of large muscularized vessels in the mutant mice. The error bars are reported as SEM. **B)** Representative images of the ROSA26-BMPR2^R899X^ mutants and wild type mice illustrate indistinguishable changes in muscularized vessels after administration of saracatinib and dasatinib. Images are taken at 10x.

## Discussion

These results suggest that saracatinib and dasatinib do not prevent the onset of PAH despite inhibiting the downstream activity of phosphorylated SRC due to the BMPR2 mutation (**Figure 1**). Wild type BMPR2’s cytoplasmic tail binds with SRC but does not lead to its abnormal phosphorylation. The truncated cytoplasmic tail in the mutant BMPR2 protein results in both the increase in phosphorylation and downstream activity of SRC. The administration of saracatinib and dasatinib are known to inhibit SRC’s activity (15, 16). Previous studies have shown that antagonizing other proteins in the SRC pathways have resulted in the amelioration of PAH in identical animal models (14). Here, we show that inhibiting this downstream activity by other therapeutic agents does not prevent the development of PAH.

Western blots confirm that both saracatinib and dasatinib inhibit as expected, with decreased expression of a key protein downstream of SRC. The animal model also revealed an expected penetrance and development of PAH, as measured through RVSP **(Figure 2).** Despite therapeutic intervention, the elevated RVSP in the Rosa26-Bmpr2^R899X^ mutant mice was not lowered. This finding suggests that the inhibition of SRC and its downstream activity do not reduce the elevated pressures that are hallmark of PAH.

Rosa26-Bmpr2^R899X^ mice had about twice the amount of large fully muscularized vessels when compared to the wild type mice. The administration of saracatinib and dasatinib failed to normalize the number of vessels. This finding once again suggests that saracatinib and dasatinib do not prohibit the development of heritable PAH.

Because previous success was seen by antagonizing the SRC pathway in PAH, further efforts should focus on understanding the complexity of the interaction between SRC and the truncated cytoplasmic tail in the BMPR2 mutation in PAH. It is plausible that the development of PAH may not be due to the phosphorylation or downstream activity associated with SRC’s differential regulation. Other studies have shown that SRC trafficking and localization is altered within the mutant cells when compared to the wild type cells (14). A previous study interrogated the SRC pathway in heritable PAH and found success in preventing the onset of disease (14). However, a key difference is that this previous study also aimed to restrict the aberrant SRC trafficking seen in mutant cells. Therefore, it is possible that the differential trafficking of SRC in heritable PAH plays a significant role in disease development and the inhibition of SRC itself is only part of the strategy that would need to be adopted the treat the disease. SRC is known to be sequestered in its phosphorylated form in PAH, and is thus unable to activate distant targets. This sequestration may be the dominating problem in SRC’s role in PAH, and inhibiting SRC’s phosphorylation may be the incorrect target to prevent heritable PAH (14). Interestingly, one of the therapies, dasatinib, is known to cause PAH in humans (19). This clinical observation further bolsters our belief that inhibiting SRC does not prevent disease, and the localization and trafficking is a more likely player in pathogenesis. Our study indicates that more information is needed to understand the complex role of SRC and the development of PAH. The inhibition and normalization of SRC’s phorpshorylation and downstream target activity alone is not sufficient to prevent heritable PAH.

## Acknowledgements

None

## Non-standard Abbreviations and Acronyms

αSMA: smooth muscle α-actin
CAS: p130Cas
BMP: bone morphogenetic protein
BMRP2: bone morphogenetic protein type II receptor
PAH: pulmonary arterial hypertension
RVSP: right ventricular systolic pressure
RTK: receptor tyrosine kinases
SFK: Src family kinases

